# Preventing extinctions post-2020 requires recovery actions and transformative change

**DOI:** 10.1101/2020.11.09.374314

**Authors:** Friederike C. Bolam, Jorge Ahumada, H. Reşit Akçakaya, Thomas M. Brooks, Wendy Elliott, Sean Hoban, Louise Mair, David Mallon, Philip J.K. McGowan, Domitilla Raimondo, Jon Paul Rodríguez, Dilys Roe, Mary B. Seddon, Xiaoli Shen, Simon N. Stuart, James E.M. Watson, Stuart H.M. Butchart

## Abstract

Stopping human-induced extinctions will require strong policy commitments that comprehensively address threats to species. In 2021, a new Global Biodiversity Framework will be agreed by the Convention on Biological Diversity. Here we investigate how the suggested targets could contribute to reducing threats to threatened vertebrates, invertebrates, and plants, and assess the importance of a proposed target to implement recovery actions for threatened species. We find that whilst many of the targets benefit species, extinction risk for over one third of threatened species would not be reduced sufficiently without a target on recovery actions, including *ex situ* conservation, reintroductions and other species-specific interventions. A median of 41 threatened species per country require such actions, and they are found in most countries of the world. To prevent future extinctions, policy commitments must include recovery actions for the most threatened species in addition to broader transformative change.

## Introduction

The world is facing an extinction crisis, with over 32,000 species documented as threatened (IUCN 2020), and extrapolations indicating that one million species are at risk of extinction (Díaz et al., 2019). Halting extinctions and reducing extinction risk is addressed in the UN Sustainable Development Goals, where Target 15.5 commits governments to *“by 2020, protect and prevent the extinction of threatened species”.* A key policy mechanism to reverse species loss is the Convention on Biological Diversity (CBD) to which 195 national governments are party.

In 2021, the Parties to the CBD will adopt a new Global Biodiversity Framework. The latest draft, published in August 2020, includes four goals and 20 targets to achieve the four goals (Secretariat of the CBD, 2020a). Goal A would commit countries to improving the status of natural ecosystems and *“reducing the number of species that are threatened by [X%]”* and maintaining genetic diversity by 2050. Rounsevell et al. (2020) suggest that reducing extinction rates should be an overarching target for the CBD, analogous to the 2°C climate target, and emphasising the importance of saving species. The CBD’s previous target to prevent extinctions and improve the status of threatened species by 2020 was not achieved (Secretariat of the CBD, 2020b). While some extinctions were prevented (Bolam et al. 2020), other species were lost, including Pinta Giant Tortoise *Chelonoidis abingdonii* and Alagoas Foliage-gleaner *Philydor novaesi* (IUCN 2020). A total of 23.7% of species remain listed as threatened with extinction of those taxonomic groups that have been comprehensively assessed on the IUCN Red List (Secretariat of the CBD, 2020b). On average, vertebrate populations are estimated to have declined (Inger et al. 2014, WWF 2020.)

The post-2020 Global Biodiversity Framework will set the global conservation agenda for the next decade. To learn from the past and avoid future human-induced extinctions, it is important to evaluate whether the proposed targets will be adequate for halting the extinction of threatened species. We assess how individual targets potentially contribute to reducing threats to species. We identify how many species would benefit from targets that address major drivers of species loss. We also identify those species that will remain threatened without a target on species-specific recovery actions, because the threats to their survival are not addressed by the other targets or because species-specific recovery actions have been identified as critical for their survival.

## Methods

We considered seven of the 20 proposed targets (Secretariat of the CBD, 2020a) - those that address threats to biodiversity and active species management. The seven targets aim to (1) implement spatial planning to retain and restore ecosystems and connectivity, (2) protect and conserve sites of particular importance for biodiversity, (3) ensure active management to enable species recovery, and reduce human-wildlife conflict, (4) ensure harvesting, trade, and use of species is legal and sustainable, (5) address invasive species, (6) reduce pollution, and (7) contribute to climate change mitigation and adaptation. To identify the number of threatened or Extinct in the Wild species that would benefit from each of the proposed targets, we matched threats to species with those that we judged would be addressed by each target. We treated Target 3 differently as it is not about a particular threat, but to encourage active species management.

### Taxonomic groups included

We downloaded IUCN Red List of Threatened Species (IUCN 2020, hereafter Red List) information for all comprehensively assessed taxonomic groups at a global level (36,602 species) on 12 May 2020, and retained all species listed as threatened (i.e. in the Red List categories of Critically Endangered, Endangered or Vulnerable) or Extinct in the Wild (7,313 species): amphibians (2,204 species), birds (1,491), mammals (1,248), selected dicot groups (683), selected crustacean groups (482), reefforming corals (232), sharks, rays and chimeras (206), conifers (205), selected bony fishes groups (202), cycads (196), selected reptile groups (100), selected gastropod groups (41), hagfish (9), cephalopods (5), gnetopsida (4), coelacanths and lungfish (3), and horseshoe crabs (2).

### Matching threats to targets

Pressures on species are documented on the Red List using hierarchical classification schemes for threats and stresses (Salafsky et al., 2008). The threats are grouped into 12 broad categories, including biological resource use, pollution, and climate change and severe weather. Each record of a threat to a species also has corresponding stresses listed (i.e. how the threat is affecting the species, for example through ecosystem degradation or species mortality). Of species we considered, 98% have at least one threat listed in their assessments. The threats have corresponding stresses listed for 97% of species-threat records.

We matched each threat-stress combination to the proposed targets (see supplementary material) because different stresses resulting from each threat may be addressed by different targets. We excluded natural threats such as volcanoes and earthquakes, which cannot be easily mitigated and are documented as threatening only 141 species. We grouped Target 1 (Spatial planning to retain and restore ecosystems) and Target 2 (Protect sites for particular importance for biodiversity) as their impacts on threats and stresses to species cannot be disentangled. We then calculated the number of species affected by each threat-stress combination. Because documentation of stresses and conservation actions needed on the Red List may not be comprehensive, it is possible that the findings presented here underestimate the number of species that would benefit from achievement of each target.

### Identifying species needing recovery actions

To identify species that would benefit from the proposed Target 3 (Ensure active management to enable species recovery), we first identified species that are affected by threats not addressed by any of the other targets. We then added those species that require species-specific conservation actions as listed on the Red List (species recovery, species reintroduction, and *ex situ* conservation). This information was available for 84% of the threatened species we analysed. Using data from the Red List, we mapped the distribution at country-level for species that require Target 3.

We also identified species with very small population sizes, making them highly susceptible to inbreeding depression, allee effects (inability to find mates), lack of genetic variation for adaptation, and stochastic events. Such species may not fully recover without the measures proposed in Target 3. Specifically, we identified species with a minimum population size below 1,000 mature individuals, those assessed under Red List criterion D or D1, those assessed as Critically Endangered under criterion C, or Endangered or Vulnerable under criterion C2ai. These criteria are triggered if the number of mature individuals, or the number in each subpopulation, is below 1,000. We also included species with severely fragmented ranges and extreme fluctuations (criterion Bac).

## Results

There are substantial differences in the number of species that would benefit from each target, according to the threats coded for each species (Fig. 1A). Target 1 (Using spatial planning to retain and restore ecosystems) and Target 2 (Protect and conserve sites for particular importance for biodiversity) combined will be particularly important as 83% of threatened and Extinct in the Wild species (6,058 species) would benefit from their implementation. This is followed by Target 4 (Ensure harvesting, trade and use of wild species is legal and at sustainable levels) with 63% (4,596 species), Target 5 (Address invasive species) with 23% (1,695 species), Target 6 (Reduce pollution) with 20% (1,472 species) and Target 7 (Climate change mitigation and adaptation) with 18% (1,339 species).

**Figure 1.**
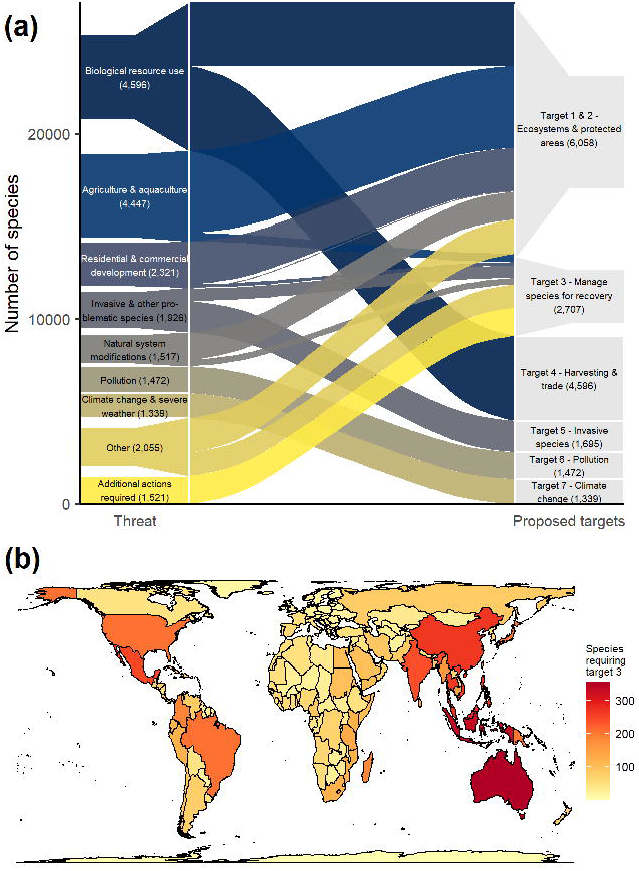
A. Number of threatened and Extinct in the Wild species whose threats are addressed by the proposed post-2020 targets for all comprehensively assessed species groups on the IUCN Red List. Species (N = 7,313) can be affected by more than one threat, and threats can be tackled by more than one target. Colours distinguish different threats. Threats are based on the IUCN Red List classification, except for *additional actions required* (see methods). B. Number of species per country that require implementation of Target 3.

At least 37% of threatened and Extinct in the Wild species (2,707 species) would likely require Target 3 (Ensure active management to enable species recovery) (Fig. 1A). These comprise 1,977 species that are affected by threats not addressed in the proposed targets, and 1,521 species that need species recovery actions, *ex situ* conservation, and/or reintroductions (with an overlap of 791 species). Species potentially requiring Target 3 occur in almost every country of the world, with a median of 41 species per country (fig. 1B). Australia supports most species (356), followed by Indonesia (334) and Malaysia (278). Additionally, a further 489 species have population sizes below 1,000 and may also benefit from Target 3.

Some actions necessary for conserving threatened species according to the Red List are addressed by the proposed post-2020 action targets that focus on mitigating threats, such as site and area protection and management, necessary for 5,053 species (Fig. 2). Most of such actions would however only be covered under Target 3, such as *ex situ* conservation (listed for 1,142 species), species recovery actions including vaccinations, supplementary feeding, or breeding site provision (681 species), and species re-introductions (260 species).

**Figure 2.**
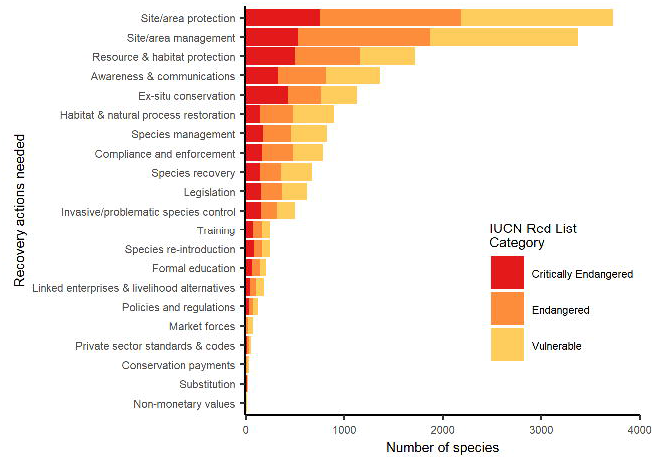
Number of threatened species that need different types of conservation actions, as identified through the IUCN Red List (IUCN 2020), by IUCN Red List category. The 15 species listed as Extinct in the Wild were excluded as there are too few to visualise in this figure.

## Discussion

Our analysis provides an indication of the relative importance of different targets for achieving the goal for conserving threatened species. Maintaining ecosystems and protected areas will play a key role, since 83% of threatened and Extinct in the Wild species could benefit from them. Other key actions include managing unsustainable harvesting and trade (addressed by Target 4, 63% of species), and controlling invasive species (Target 5, 23% of species). However, Target 3 will be essential in promoting the recovery of over one in three threatened and Extinct in the Wild species, because their threats are not addressed by the other targets, or because they require targeted species-specific actions. Our results emphasise how critical it is to retain such a target in further negotiations.

### Tackling the most pervasive threats

The CBD’s post-2020 Global Biodiversity Framework needs to lead to the transformative change required for halting species extinctions (Díaz et al., 2020), by addressing the underlying drivers of species loss. Tackling threats is important for currently threatened species, but also for preventing even more species from becoming threatened. Our results highlight the importance of targets that aim to tackle the most pervasive threats to species, particularly land use change through agriculture and overexploitation. There are transformative pathways that show we can maintain ecosystems whilst ensuring food security, by making food production more sustainable, changing consumption and diet choices to sustainable and healthy levels, and increasing protected area coverage (Leclère et al., 2020), all of which are consistent with the draft targets.

To ensure the proposed targets will lead to halting extinctions however, two further assumptions must be met: that targets address threats sufficiently to reduce extinction risk, and that targets are fully and effectively implemented (Díaz et al., 2020). For example, threatened species need adequate representation in the network of protected areas and other effective area-based conservation measures, by securing sites such as KBAs that are critical in their conservation value (Visconti et al., 2019). Such species not only need sufficient coverage by protected and conserved areas, but also that these are effectively and equitably managed and appropriately connected (Maxwell et al., 2020). While effective management and connectivity are part of the draft Target 2 wording, equitable management is not, even though it is known to lead to better outcomes for both people and nature (Oldekop et al., 2016), and is in line with some of the other draft targets.

### Species that require recovery actions to ensure their survival

Our analysis has demonstrated that in order to achieve Goal A, it is essential to retain Target 3 in the Post-2020 Global Biodiversity Framework, to ensure active management to enable species recovery. Target 3 will be necessary for 2,707 species that are facing threats not tackled by other targets, or that will require species-specific recovery actions. Examples include 238 endemic Hawaiian plant species with fewer than 50 individuals remaining in the wild (Werden et al., 2020), such as the Punaluu Haha *Cyanea truncata* which requires intensive *in situ* recovery actions to manage the threat of invasive species as well as *ex situ* conservation to supplement the population. For other plant species, labour-intensive planting, watering and protection of seedlings is needed due to no natural regeneration, such as the iconic oak *Quercus brandegeei* in Mexico (Denvir et al., 2016), and the Baishan Fir *Abies beshanzuensis* in China (Yang et al., 2013). The Lord Howe Island Stick-insect *Dryococelus australis* has no more than 35 surviving individuals in the wild, but once invasive plants are removed from its range on a small island, re-introduction efforts will take place using individuals from *ex situ* populations in zoos that number in the thousands (Rudolph and Brock, 2017).

For other species, we do not yet fully understand how to tackle the threats they face, such as 232 threatened coral species impacted by bleaching, 571 threatened amphibian species impacted by chytridimycosis, or those species whose mutualists (seed dispersers, pollinators, symbionts) have disappeared locally or globally. For such species, *ex situ* conservation may ‘buy time’ while feasible interventions are devised, tested, and applied (da Silva et al., 2019). This would ensure that species can be re-introduced, or populations supplemented.

There is evidence that we can prevent extinctions even of those species at the brink of extinction (Bolam et al., 2020). For a subset of threatened species, these actions are not only necessary but also achievable if there is political will and resources available to reverse declines. There are examples of species that have recovered rapidly owing to recovery actions, such as the Seychelles Warbler *Acrocephalus sechellensis* which was listed as threatened in 1988 and had recovered to Near Threatened by 2015 due to translocations and habitat management (BirdLife International, 2016). To prevent further extinctions, these actions need to be underpinned by strong policy commitments so they can be scaled up.

### A target for species recovery actions post-2020

Our results demonstrate the importance of retaining Target 3 in future negotiations to prevent further extinctions. The current wording, *“By 2030, ensure active management actions to enable wild species of fauna and flora recovery and conservation, and reduce human-wildlife conflict by [X%]”,* would benefit from greater detail, for example, *“Implement intensive species-specific recovery actions by 2030, in situ and ex situ, where required, for species whose survival depends on such actions or whose recovery cannot otherwise be enabled or sustained.”* We also suggest that the need to address human-wildlife conflict would be more appropriately included in draft Target 4 on harvesting, trade and use of species, rather than in Target 3.

If sufficiently implemented, our proposed target wording would contribute to achieving the 2050 draft goal of reducing the number of species that are threatened. Target 3 could be monitored using indicators based on the IUCN Red List, including the Red List Index (measuring trends in extinction risk for sets of species, Butchart et al., 2004, Butchart et al., 2007). It could be informed by the establishment of science-based targets for species using the Species Threat Abatement and Restoration metric (Mair et al. in review) and by Green Status of Species assessments (Akçakaya et al., 2018).

The draft targets of the post-2020 Global Biodiversity Framework cover the key threats to species. In addition, Target 3 covers the interventions required for those species in need of additional recovery actions. Therefore it is critical that all draft targets are retained in the final framework. Further human-induced species extinctions can be prevented, but only if both threats to species are addressed and species recovery actions are implemented as a matter of urgency.

## Acknowledgments

We thank Craig Hilton-Taylor for provision of data and Bob Lacy for comments on an early draft.

## Author contributions

Conceptualization: FCB, TMB, LM, PJKM, SNS, JEMW, SHMB; Methodology: FCB, HRA, TMB, LM, SHMB; Formal analysis: FCB; Data curation: FCB; Writing - original draft: FCB, TMB, SHMB; Writing - reviews& editing: FCB, JA, HRA, TMB, WE, SH, LM, DM, PJKM, DRa, JPR, DRo, MBS, XS, SNS, JEMW, SHMB; Visualization: FCB; Supervision: PJKM, SHMB; Project administration: FCB, PJKM.

## Data and materials availability

Data and code for reproducing numbers and figures shown are available at https://github.com/rbolam/Species_target.

